# DNA sequence+shape kernel enables alignment-free modeling of transcription factor binding

**DOI:** 10.1101/089441

**Authors:** Wenxiu Ma, Lin Yang, Remo Rohs, William Stafford Noble

## Abstract

**Motivation:** Transcription factors (TFs) bind to specific DNA sequence motifs. Several lines of evidence suggest that TF-DNA binding is mediated in part by properties of the local DNA shape: the width of the minor groove, the relative orientations of adjacent base pairs, etc. Several methods have been developed to jointly account for DNA sequence and shape properties in predicting TF binding affinity. However, a limitation of these methods is that they typically require a training set of aligned TF binding sites.

**Results:** We describe a sequence+shape kernel that leverages DNA sequence and shape information to better understand protein-DNA binding preference and affinity. This kernel extends an existing class of *k*-mer based sequence kernels, based on the recently described di-mismatch kernel. Using three *in vitro* benchmark datasets, derived from universal protein binding microarrays (uPBMs), genomic context PBMs (gcPBMs) and SELEX-seq data, we demonstrate that incorporating DNA shape information improves our ability to predict protein-DNA binding affinity. In particular, we observe that (1) the *k*-spectrum+shape model performs better than the classical *k*-spectrum kernel, particularly for small *k* values; (2) the di-mismatch kernel performs better than the *k*-mer kernel, for larger *k*; and (3) the di-mismatch+shape kernel performs better than the di-mismatch kernel for intermediate *k* values.

**Availability:** The software is available at https://bitbucket.org/wenxiu/sequence-shape.git

**Supplementary information:** Supplementary data are available at *Bioinformatics* online.

## 1 Introduction

Modeling transcription factor (TF) binding affinity and predicting TF binding sites are important for annotating and investigating the function of cis-regulatory elements. In the past decade, the development of chromatin immunoprecipitation coupled with high-throughput sequencing (ChIP-seq, Johnson *et al.*, 2007; Robertson *et al.*, 2007; Barski *et al.*, 2007), protein binding microarrays (PBMs, Berger *et al.*, 2006) and systematic evolution of ligands by exponential enrichment coupled with high-throughput sequencing (SELEX-seq, Zykovich *et al.*, 2009; Zhao *et al.*, 2009; Jolma *et al.*, 2010; Slattery *et al.*, 2011) has provided high-resolution TF binding datasets both *in vivo* and *in vitro*. However, despite the increasingly large collection of such datasets, our ability to predict where a given TF binds to genomic DNAs is still imperfect.

One important challenge associated with TF binding prediction is how to properly model combinatorial binding that involves multiple TFs or the effects of local chromatin architecture. Recent studies have shown that the interaction of the TF with co-binding factors (Lemon and Tjian, 2000; Slattery *et al.*, 2011) and local chromatin architecture (Boyle *et al.*, 2011; Dror *et al.*, 2015) affects TF binding to target sites. Hence, computational © The Author 2016. Published by Oxford University Press. All rights reserved. For permissions, please methods that explicitly model cis-regulatory modules (Zhou and Wong, 2004; Kato *et al.*, 2004) and local chromatin accessibility (Hesselberth *et al.*, 2009; Chen *et al.*, 2010) have been developed to address these issues.

However, as evidenced by our inability to predict *in vitro* binding derived from high-throughput assays such as PBMs or SELEX-seq experiments, combinatorial factors are not the only culprit. A second challenge lies in building computationally tractable, physically plausible models. For example, commonly used position weight matrix (PWM) methods depend on correctly aligned DNA sequences and make the unrealistic assumption that each nucleotide binds to the TF independently of one another. Accordingly, a variety of methods have been proposed that attempt to expand this approximation (Barash *et al.*, 2003; Zhou and Liu, 2004; Sharon *et al.*, 2008; Zhao *et al.*, 2012).

Dependencies between nucleotide positions in a TF binding site can be explicitly encoded through *k*-mers, for instance dinucleotides or trinucleotides (Zhao *et al.*, 2012; Gordân *et al.*, 2013). On the other hand, because stacking interactions between adjacent base pairs give rise to three-dimensional DNA structure, DNA shape features represent an alternative approach for encoding nucleotide dependencies implicitly (Zhou *et al.*, 2015). Recent evidence suggests that a crucial aspect of TF binding can be explained based on the DNA shape of selected targeted sites (Rohs *et al.*, 2009). Local structural features of the double helix, such as minor groove width (MGW), roll, propeller twist (ProT) and helix twist (HelT), have been proven to greatly affect TF binding (Zhou *et al.*, 2015). Therefore, whereas traditional TF binding prediction takes as input only the primary nucleotide sequence, improved performance can be obtained by taking into account aspects of the DNA shape (Zhou *et al.*, 2015; Gordân *et al.*, 2013; Levo *et al.*, 2015). This approach has the potential to significantly improve our ability to predictively model TF-DNA interactions *in vitro* (Abe *et al.*, 2015) and *in vivo* (Mathelier *et al.*, 2016).

In this study, we developed a kernel-based regression and classification framework that enables accurate and efficient modeling and prediction of TF-DNA binding affinities. One of the most compelling motivations for using kernel functions is that kernels can be defined over arbitrary types of heterogeneous objects, such as pairs of vectors, discrete strings of variable length, graphs, nodes within graphs, trees, etc. (reviewed in Schoelkopf *et al.*, 2004). In our task, we used kernel functions to measure similarity between DNA sequences and between local DNA shape features, simultaneously. We propose two shape-augmented kernel functions. One is the spectrum+shape kernel (Section 2.2), which is a natural extension of the classic *k*-mer spectrum kernel (Leslie *et al.*, 2002). The other is a di-mismatch+shape kernel (Section 2.4), which is built upon the recently developed di-mismatch kernel (Agius *et al.*, 2010; Arvey *et al.*, 2012) and encodes both nucleotide sequence degeneracy and DNA shape readout.

We used these kernels in regression models, applied to both universal PBM (uPBM) and genomic-context PBM (gcPBM) data derived from a large collection of human and mouse TFs (Zhou *et al.*, 2015). Our results suggest that adding shape information substantially improved our TF binding prediction accuracies. Furthermore, we applied our di-mismatch+shape kernel in a classification setting and successfully distinguished binding sites of two homologous Hox TFs using SELEX-seq data (Abe *et al.*, 2015). We thus found that our shape-augmented model accurately detected subtle but important differences in local DNA shape conformations.

## 2 Approach — Kernel Methods

In this study we devised and evaluated several kernel methods for building quantitative models of TF binding affinity. In each case, we consider the following problem. Suppose we are given a collection of triples (*q*_1_, *x*_1_, *y*_1_), …, (*q_n_, x_n_, y_n_*), where *q_i_* is a DNA sequence of length *w*, *x_i_* contains information about the DNA shape conformation of *q_i_*, and *y_i_* is either a real number that indicates the relative strength of binding of a particular TF to *q_i_* (in a regression setting) or a binary indicator that the TF either binds to the sequence or does not bind (in a classification setting). Our goal is to build a predictive model *f*(.) such that *f*(*q_i_, x_i_*) = *y_i_*. We consider a variety of kernel methods for projecting either *q_i_* or *q_i_* and *x_i_* into a vector space suitable for a classical regression or classification algorithm.

### 2.1 Spectrum kernel

A simple and widely used kernel for representing biological sequences is the *spectrum kernel* (Leslie *et al.*, 2002). This kernel is defined over an *n*-dimensional feature space, where *n* is the number of unique *k*-mers in the dataset. Note that, due to the reverse complementarity of DNA sequences, *n* = 4*^k^* /2 if *k* is odd and *n* = (4*^k^* + 4*^k^* /2)/2 otherwise (Supplementary Table 1). Each feature corresponds to a unique string of length *k*, and the feature values are counts of the number of times the given string occurs within the given DNA sequence. The kernel is a scalar product in this feature space, which can be computed efficiently using several different data structures (Leslie *et al.*, 2002; Vishwanathan and Smola, 2003). The hyperparameter *k* determines the dimensionality of the feature space. An important characteristic of the spectrum kernel is that it is compositional rather than positional; i.e., the position of the *k*-mer within the given sequence has no effect on the embedding. The spectrum kernel was originally described for protein homology detection (Leslie *et al.*, 2002), but has been used for a variety of DNA-based classification and regression tasks, including predicting nucleosome positioning (Peckham *et al.*, 2007) and splice site prediction (Sonnenburg *et al.*, 2007).

### 2.2 Spectrum+shape kernel

Because we know that TF binding is mediated in part by the shape of the DNA binding site, we incorporated local DNA shape properties into our prediction models. Specifically, we considered four DNA shape features: MGW, Roll, ProT and HelT. These features were derived from Monte Carlo simulations using a previously described pentamer model (Zhou *et al.*, 2013; Chiu *et al.*, 2016). The MGW and ProT features are defined at each nucleotide position, whereas Roll and HelT define translations and rotations between two adjacent nucleotides. Thus, a pentamer contributes one MGW value and one ProT value at the central nucleotide and two Roll values and two HelT values at the two central dinucleotide pairs.

To incorporate DNA shape information into the spectrum kernel, we developed a *spectrum+shape kernel*. This kernel is defined over a (3 + 4*k*) *. n*-dimensional feature space (Supplementary Table 1). The first *n* features are defined over the *n* unique *k*-mer sequences in the same manner as described for the classic spectrum *k*-mer kernel. The remaining features capture the four corresponding shape properties. Consider MGW as an example. For each unique *k*-mer, we find all its occurrences within the given DNA sequence, and we extract the *k*-mer sequences plus 2 bp flanking sequences on both sides. If the *k*-mer appears in the beginning or at the end of the given DNA sequence, then we add “NN” to its 5’ and 3’ flanks to make it of length *k* + 4. Then we calculate the average MGW values over all the extracted substrings of length *k* + 4. Since each pentamer contributes one MGW value, each (*k* + 4)-mer will contribute *k* MGW values. Therefore, we have a total of *kn* features defined for MGW shape information. In this way, we can define *kn* features each for MGW and ProT, and (*k* + 1) *. n* features each for Roll and HelT.

Note that our spectrum+shape kernel differs from the sequence+shape model used in Zhou *et al.* (2015). Our model is compositional and hence can be applied to unaligned DNA sequences. The Zhou model, in contrast, is positional and hence requires pre-alignment of the TF binding sites. This requirement used in our previous studies (Zhou *et al.*, 2015; Abe *et al.*, 2015) represents a limitation that restricted us from analyzing data that could not be aligned. Overcoming this limitation is particularly important for low affinity TF binding (Crocker *et al.*, 2015) or binding site sampling during the search process (Dror *et al.*, 2016).

**Fig. 1.**
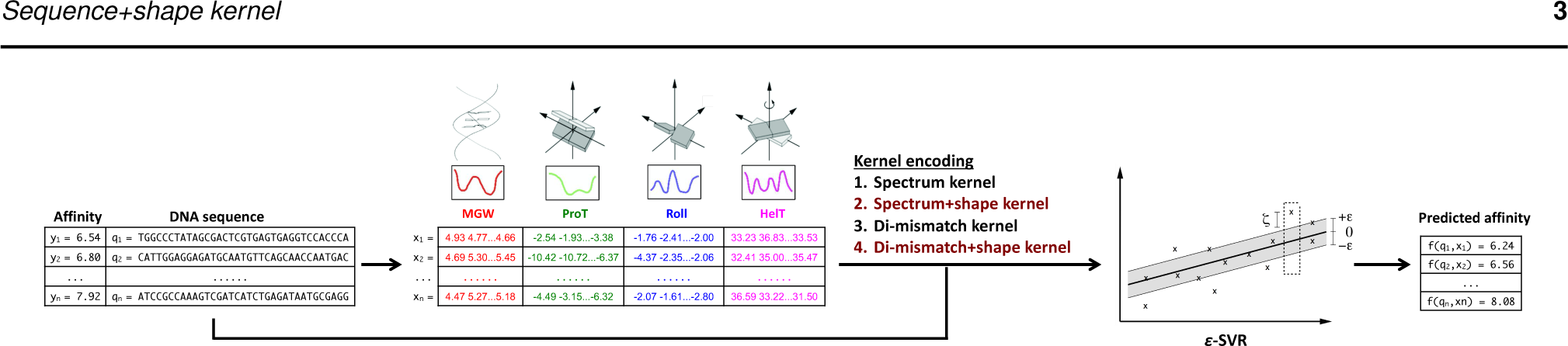
*ϵ*-Support Vector Regression (SVR) framework for the alignment-free modeling of transcription factor binding.

### 2.3 Di-mismatch kernel

Subsequent to the spectrum kernel, a variety of more complex and more powerful DNA kernels have been developed. For example, the *mismatch kernel* generalizes the spectrum kernel by relaxing the matching function on substrings (Leslie *et al.*, 2003). In the mismatch kernel, a *k*-mer is considered to occur at a specific position within the sequence *q* if the *k*-mer matches *q* with up to *m* mismatches. A more recent alternative generalization, the *di-mismatch kernel*, uses a matching function that counts the number of matching dinucleotides in the two *k*-mers (Agius *et al.*, 2010). Like the spectrum kernel, only exact matches between dinucleotides are considered; however, a second hyper-parameter *m* specifies a threshold so that the match score is set to zero if the number of matching dinucleotides falls below *k* − *m* − 1. Precisely, we let {*ϕ_i_* }_*i* =1…*n*_ be the set of unique *k*-mers that occur in a large set of training sequences. Then, given a training sequence *q* of length *w*, we define the set of substrings of length *k* in *q* to be

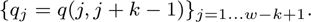

In this setting, the DNA sequence *q* may be represented by a feature vector

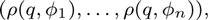

where

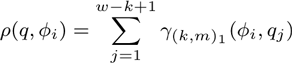

and the value *γ*(*k,m*)_1_ (*ϕ_i_*, *q_j_*) is the di-mismatch score between two *k*-mers, which counts the number of matching dinucleotides between *ϕ_i_* and *q_j_*, i.e.,

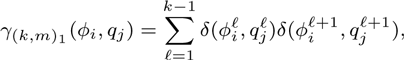

where *δ*(.) is the Kronecker delta function. The mismatch threshold *m* has the effect of setting the score to 0 if *γ*(*k,m*)_1_ (*ϕ_i_*, *q_j_*) < *k* – *m* – 1, i.e., the number of dinucleotide mismatches between *ϕ_i_* and *q_j_* exceeds *m*. This threshold forces the kernel to only consider highly similar sequences. The motivation for the di-mismatch kernel is to favor *k*-mers with consecutive mismatches over *k*-mers with non-contiguous mismatches. Previous evidence suggests that the di-mismatch kernel yields more accurate TF binding predictions both *in vitro* and *in vivo* (Agius *et al*., 2010) and helps to identify cell-type specific binding (Arvey *et al*., 2012).

### 2.4 Di-mismatch+shape kernel

We generalize the di-mismatch kernel by expanding the feature vector to include both DNA sequence and shape features:

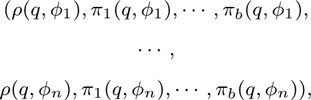

where *ρ*(*q*, *ϕ_i_*) is the previously defined di-mismatch feature function, and *π*_1_(*q*, *ϕ_i_*) to *π_b_*(*q*, *ϕ_i_*) are the DNA shape feature functions that we will introduce here.

Similar to Section 2.2, we consider the four DNA shape features: MGW, R, ProT and HelT. For each *k*-mer *ϕ_i_* (*k* ≥ 5), the sliding pentamer model (Zhou *et al.*, 2013) generates MGW and ProT feature vectors of length *k* – 4 and Roll and HelT feature vectors of length *k* – 3.

Our kernel requires that we define, for each unique *k*-mer *ϕ_i_* and *t*-th shape feature, a corresponding “canonical” shape feature vector *s_i,t_*. A simple way to define such a feature vector is by averaging over all possible 2-bp sequences immediately upstream and downstream. In this case, *s_i,t_* is a vector of length *k* for MGW and ProT, and *k* + 1 for Roll and HelT.

For each length-*k* substring *q_j_* in *q*, let *x_j,t_* be its *t*-th DNA shape feature vector, *t* = 1, 2, 3, 4. For the first and last two substrings, i.e., *j* = 1, 2, *w* – *k*, *w* – *k* + 1, *x_j,t_* can be obtained by averaging all possible 1‐ or 2-bp flanks; f or o ther i ntermediate s ubstrings, t he DNA shape features can be obtained directly.

Thus we define the *t*-th DNA shape feature function as

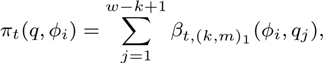

where the shape feature similarity score *β*_t_,(*k,m*)_1_ (*ϕ_i_*, *q_j_*) is

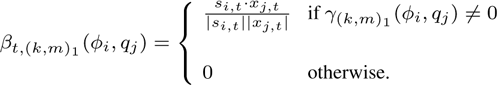

That is, the shape similarity score equals the normalized inner product between the shape feature vectors *s_i,t_* and *x_j,t_*, and we set the score to zero if the number of dinucleotide mismatches between *ϕ_i_* and *q_j_* exceeds the threshold *m*. This generalized di-mismatch kernel is defined over a 5*n*-dimensional feature space (Supplementary Table 1).

## 3 Methods

### 3.1 TF binding datasets

We used three types of *in vitro* datasets to evaluate and compare the performance of the kernels described above.

The universal PBM (uPBM) data from the DREAM5 project (Weirauch *et al.*, 2013, GEO accession number GSE42864) consists of unaligned 35-mer PBM probes for 66 TFs from a variety of protein families. The normalized uPBM data were downloaded from the DREAM5 challenge website, where the data was normalized according to the total signal intensity. Unlike in Zhou *et al.* (2015), we did not align or trim the probes based on the reported motifs of the biniding sites.

The genomic context PBM (gcPBM) data are for three human basic helix-loop-helix (bHLH) TF dimers: Mad1 (Mxd1)–Max, Max–Max, and c-Myc–Max (Mad, Max, and Myc, respectively) (Zhou *et al.*, 2015, GEO accession number GSE59845). The gcPBM data consists of 36-mer probes, each in its real genomic context. The gcPBM probes were by design pre-aligned at the center using the E-box motif sites. We used the raw probes in this study, without any filtering for absence or multiple occurence of E-box binding sites.

The homeodomain (Hox) data consists of SELEX-seq data for two *Drosophila* Hox proteins, Scr and Antp, each in complex with Exd (Abe *et al.*, 2015, GEO accession number GSE65073). These two Exd-Hox dimers bind to a similar consensus motifs but have distinct DNA shape preferences. The SELEX-seq-derived 16-mers and their TF binding affinities were obtained from Abe *et al.* (2015). No further filtering using either Hox monomer or Exd-Hox heterodimer motifs was performed. Each sequence in this dataset is associated with a relative TF binding affinity, normalized to values ranging from 0.0 to 1.0. For each sequence, we calculated separately the percentiles of relative binding affinity for the Scr and Antp bound sequences, respectively. Sequences with a relative binding affinity greater than 0.57 (median value in the Scr data) for Scr and less than 0.27 (the median value in the Antp data) for Antp were labeled as positive. Conversely, the sequences with a relative binding affinity of greater than 0.57 for Scr and less than 0.27 for Antp were labeled as negative. We used the resulting sequences and binary labels (Supplementary Figure 1) for the classification task.

### 3.2 Regression experiment design

We evaluated our models separately on each TF in each dataset. To archieve this, we randomly sampled 1000 input DNA sequences and their relative binding affinity values to evaluate our regression models, which significantly reduced the computational cost for kernel calculation and SVR learning.

We tested each kernel in the context of linear support vector regression (*ϵ*-SVR). We implemented the SVR framework with different kernels using the Python scikit-learn/svm module, which uses LIBSVM (Chang and Lin, 2001) as its internal SVR implementation.

To avoid over-fitting, we performed nested cross-validation (CV). The inner five-fold CV performs hyperparameter grid search. The grid includes the two SVR parameters, *C* and *ϵ* (*C* from -3 to 3 in log 10 space, *ϵ* =|0, 0.001, 0.01, 0.1, 0.2, 0.5, 1.0}). The outer five-fold CV evaluates the performance of the best model selected from the inner CV. We used the coefficient of determination *R*^2^ to measure the kernel performance. The *R*^2^ measurement has been used previously to evaluate regression performance for SELEX-seq and PBM data (Zhou *et al.*, 2015; Abe *et al.*, 2015). We did not use the Spearman correlation coefficient as the metric because the rank transformation results in an undesirable emphasis on the unbound, low intensity probes (Weirauch *et al.*, 2013).

To restrict the dimensionality of the feature space and improve computational efficiency, we selected the top 1,000 features for each model based on their *R*^2^ values for predicting binding affinities. To avoid over-fitting, we performed this feature selection separately in each outer CV, using the binding affinity values of the training data only.

### 3.3 Classification experiment design

We used the linear support vector machine (SVM) as our training and testing framework for the classification task.

Similar to the SVR framework, we performed nested CV to avoid over-fitting. Because of the unequal numbers of positives and negatives, we used stratified CV in both layers to equally split positive and negative labels in each fold. We used the inner five-fold CV to perform grid search for hyperparameters, which include SVM parameters (*C* in the linear SVM model, from −3 to 3 in log_10_ space). We used the outer five-fold CV to evaluate the performance of the best model selected from the inner CV. In these classification experiments, we used the area under the ROC curve (AUROC) to measure the performance. As *k* increases, the number of features increases exponentially. To restrict the dimension of the feature space and improve computational efficiency, we selected the top 1000 features for each model based on their individual AUROC scores for distinguishing between sequences with positive and negative labels. As described above, this feature selection was performed separately in each fold of the outer CV.

## 4 Results

### 4.1 In uPBM data, adding shape features yields improved performance for small *k*

To evaluate our models, we started with the uPBM datasets from the DREAM5 experiment (Weirauch *et al.*, 2013), consisting of 66 mouse TFs from various TF families. For each TF, we first evaluated the *k*-spectrum model and the *k*-spectrum+shape model, for every *k* value from 1 to 7, comparing the *R*^2^ values between the true binding affinities and our predictions.

We observed in Figure 2 that, for small *k* values (*k ≤* 5), adding shape information to the kernel leads to significantly better performance for more than 90% of the TFs (Wilcoxon’s one-sided test, *k* =1, *p* =2.07e−10; *k* =2, *p* =2.87e−7; *k* =3, *p* =5.67e−5; *k* =4, *p* =1.40e−3; *k* =5, *p* =4.93e−3). This result agrees with previous reports by Zhou *et al.* (2015). The DNA shape information is calculated based on pentamers, and therefore captures dependencies that may not be well represented by small *k*-mers. Conversely, we observed that for larger *k*, the *k*-spectrum+shape model under-performs the *k*-spectrum model. Especially when *k* > 5, the *k*-spectrum+shape model has larger variability in its performance and in some cases even yields negative *R*^2^ values. The lack of improvement from the shape features for large values of *k* is likely because the longer *k*-mers in the *k*-spectrum kernel already implicitly capture DNA shape information. Furthermore, especially for large values of *k*, shape-augmented kernels map the input sequences to a very high-dimensional feature space in which the learning task is considerably more difficult.

**Fig. 2.**
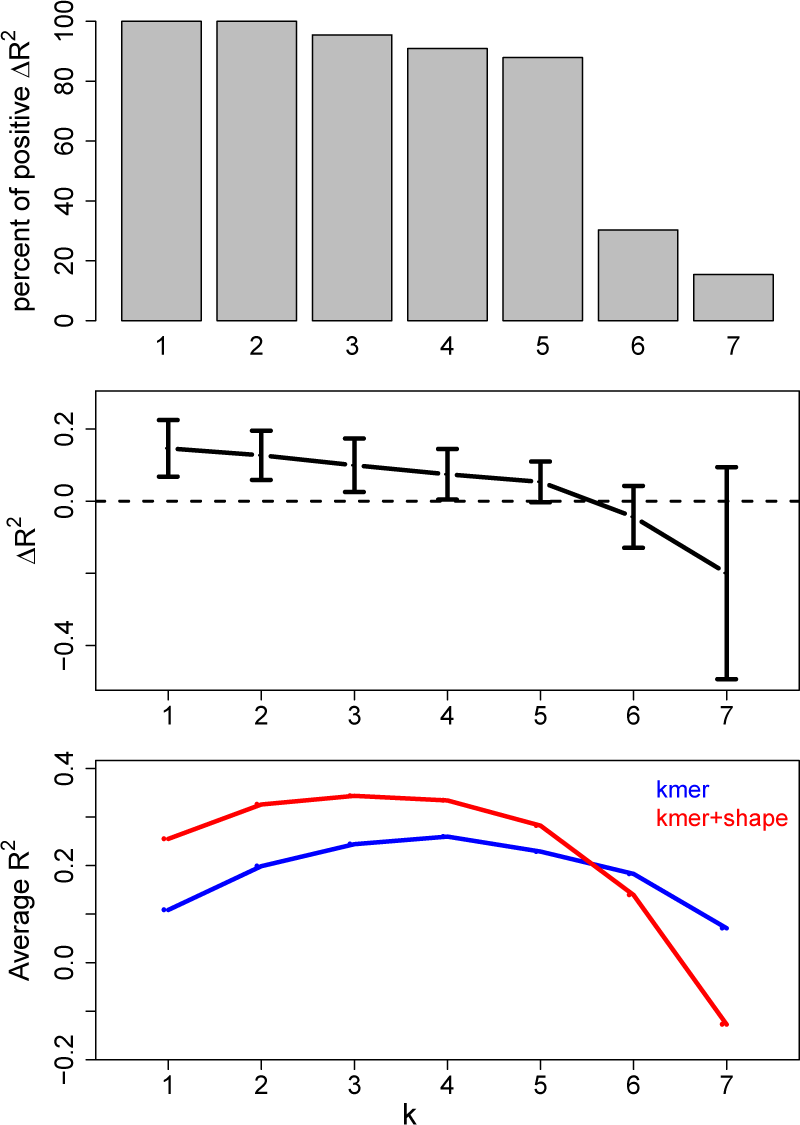
Comparison between *k*-spectrum and *k*-spectrum+shape models on uPBM dataset. The top panel is the percent of DREAM5 TFs that have higher *R*_2_ values using the *k*-spectrum+shape model than using the *k*-spectrum model; the middle panel shows the differences of *R*^2^ values between the two models; and the bottom panel shows the *R*_2_ performance scores of various *k*-spectrum models and *k*-spectrum+shape models, for *k* = 1, …, 7.

In addition to the aggregated performance over all 66 TFs, we also looked at the *R*^2^ improvement for each TF and for each TF family (Figure 3, Supplementary Figure 2). Taking *k* = 4 as an example, we found that the 4mer+shape model led to great improvements for all zinc fingers, bHLH, bZip, and helix-turn-helix (HTH) TFs. These observations are consistent with previous findings (Zhou *et al.*, 2015; Gordân *et al.*, 2013; Yang *et al.*, 2014; Stella *et al.*, 2010). Only for zinc fingers, previous studies did not detect a significant improvement in binding specificity predictions upon the addition of shape information (Zhou *et al.*, 2015). Zinc fingers recognize DNA in a modular manner with each finger binding to 3 bp, so that alignment of such modular sites is more ambiguous. The use of an alignment-free approach probes the effect of shape without the uncertainty in aligning such modular binding sites.

**Fig. 3.**
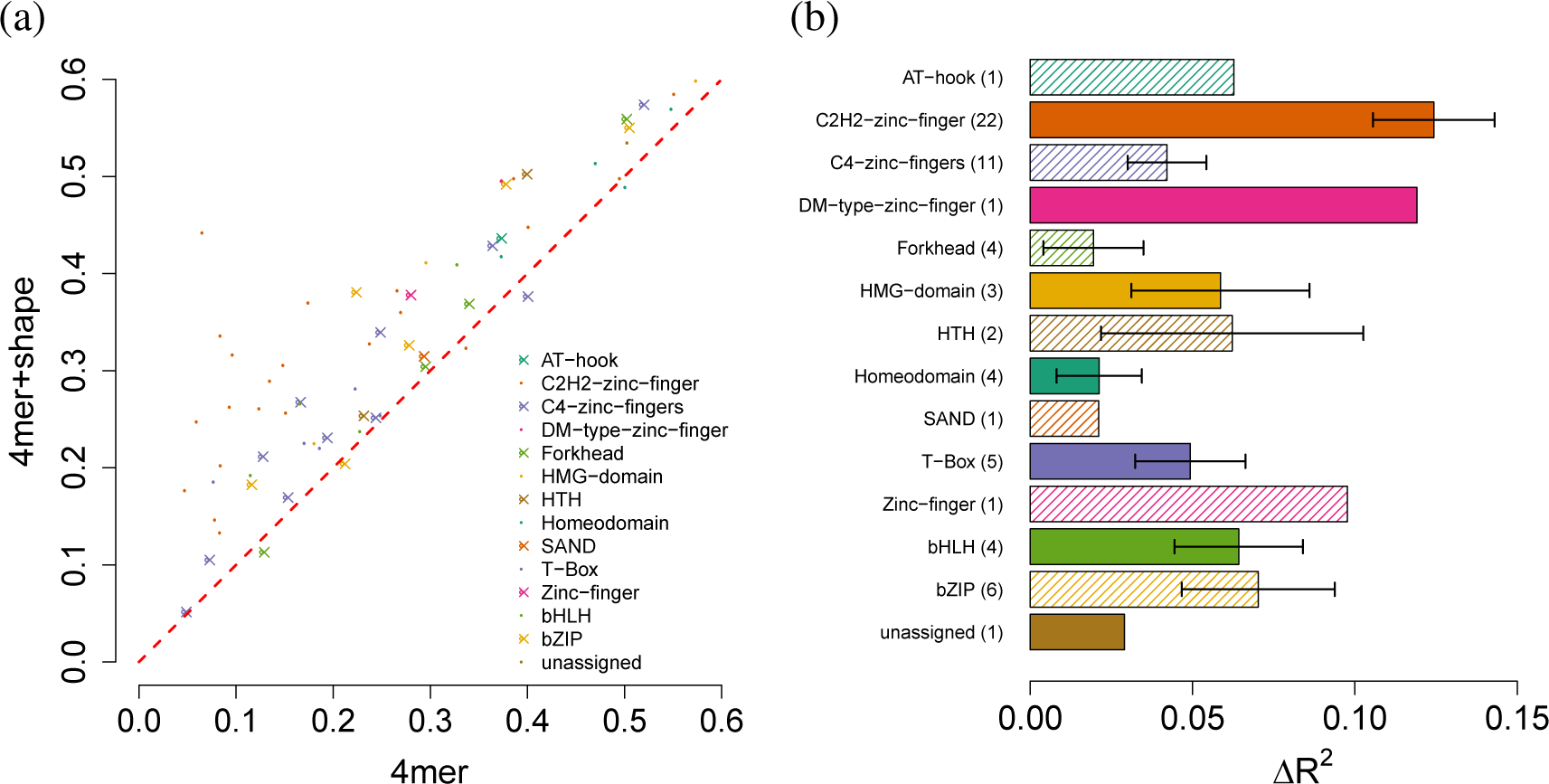
*R*^2^ performance for *k*-spectrum model verse *k*-spectrum+shape model on uPBM dataset, *k* = 4. (a) Scatter plot of the *R*^2^ performance values between the two models. Each dot represents one TF, colored corresponding to its protein family. **(b)** Bar plot of *R*^2^ improvements for various protein families. Numbers in the parentheses are the number of DREAM5 TFs in each TF family. The *x*-axis shows the differences of *R*_2_ values between the two models. The length of the bars represent the mean of *R*_2_ differences and the error bars mark the standard error of the mean.

The spectrum+shape kernel implemented in this study encodes both sequence and shape information in a compositional fashion, i.e., without respect to the absolute position of the sequence or shape feature within a given sequence. In contrast, Zhou *et al.* (2015) implemented a positional sequence+shape kernel where the input sequences are required to be aligned at the binding motif sites. In both compositional and positional models, combining sequence information and shape information contributes to the improvement of prediction performance compared to using sequence information alone. The advantage of our compositional approach is that it does not require the uPBM probes to be aligned in a pre-processing step. Taken together, the results from Zhou *et al.* (2015) and our study confirmed that DNA shape readout plays an important role in guiding TFs to recognize their target binding sites.

### 4.2 The di-mismatch kernel benefits from inclusion of shape features on uPBM datasets

Next, we compared our new di-mismatch+shape kernel with the di-mismatch kernel developed by Agius *et al.* (2010), to examine whether adding shape information to the di-mismatch kernel improves the prediction accuracy of TF binding affinities. We first implemented the di-mismatch kernel in our SVR framework and compared its performance with the spectrum kernel using the 66 mouse TFs from the DREAM5 data. In agreement with previous findings (Agius *et al.*, 2010; Arvey *et al.*, 2012), the di-mismatch kernel consistently performs better than the spectrum kernel on the uPBM DREAM5 data for large *k* values (*k* ≥ 5, Supplementary Figure 3).

We then evaluated in detail (Figure 4) at the comparison between the di-mismatch kernel with and without inclusion of shape for different *k* and *m* parameter settings (*k* = 3, …, 8 and *m* = 1, …, max|2, (*k* – 3)}). We observed several trends. First, we considered the case when we only observed one di-mismatch, i.e., *m* = 1. By definition, this can only happen when a single nucleotide mismatch occurs at the beginning or end of the *k*-mer sequence, since otherwise a single mismatch in the middle of the sequence leads to two di-mismatches. In this case, adding shape features leads to significantly improved *R*^2^ values for *k* = 3 and 4, for the majority of the TFs (Wilcoxon’s one-sided test, *k* =3, *p* =1.73e-6; *k* =4, *p* =0.04) and marginal improvements for *k* = 5 and 6 (*k* =5, *p* =0.36; *k* =6, *p* =0.35). Second, we looked at the case *m* = 2, where we allow a single mismatch to occur in the middle of the *k*-mer sequences. In that case, our di-mismatch+shape kernel performs substantialy better than the di-mismatch kernel for *k* = 4 and 5 (Wilcoxon’s one-sided test, *k* =4, *p* =2.10e−6; *k* =5,*p* =0.12). However, for *k* ≥ 6, the performance of the di-mismatch+shape model was affected by the high dimensionality of the feature space and led to worse *R*^2^ values compared to the di-mismatch model.

**Fig. 4.**
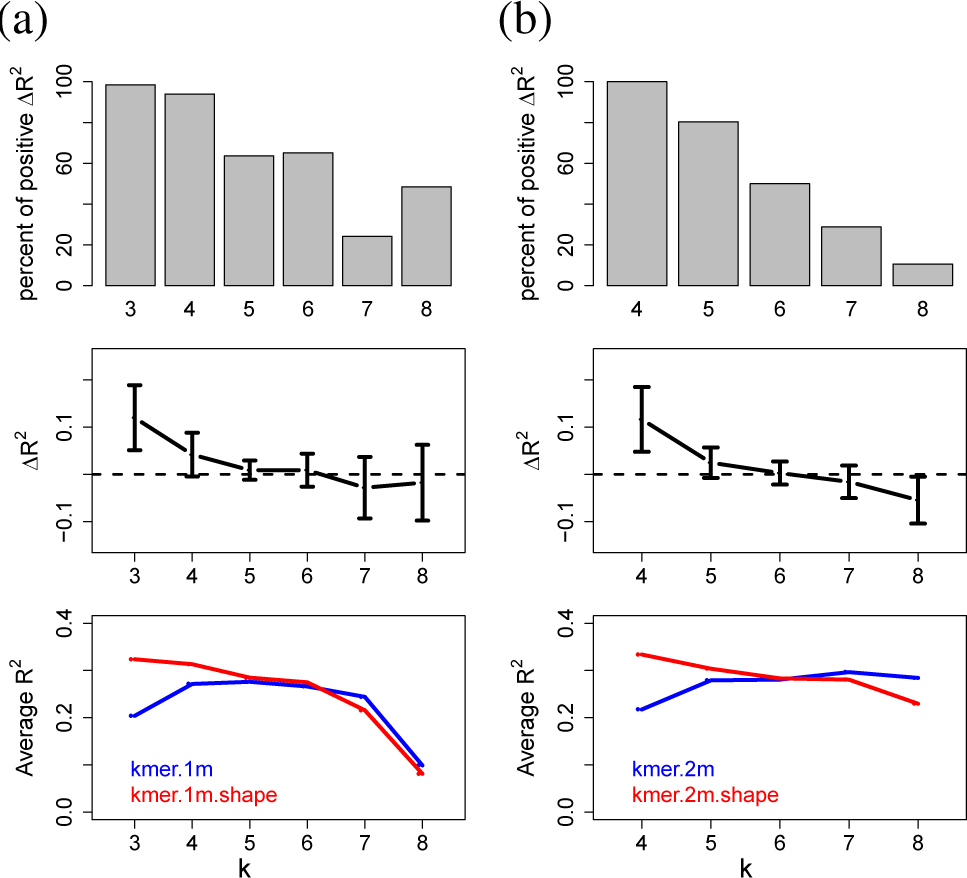
Comparison between di-mismatch and di-mismatch+shape models on uPBM dataset. **(a)** di-mismatch model verse di-mismatch+shape model for di-mismatch parameter *m* = 1, where *k* = 3, …, 8; **(b)** di-mismatch model verse di-mismatch+shape model for di-mismatch parameter *m* = 2, where *k* = 4, …, 8. The top panels are the percent of DREAM5 TF datasets that have higher *R*^2^ values using the *k*-spectrum+shape model than using the *k*-spectrum model; the middle panels show the differences of *R*^2^ values between the two models; and the bottom panels show the *R*^2^ performance scores of various di-mismatch models vs. di-mismatch+shape models.

Even though the di-mismatch kernel itself is able to encode sequence degeneracy in TF binding patterns, our results suggest that adding pentamer-based shape information to the di-mismatch kernel provides additional information about sequence dependencies and shape features, hence leading to better performance for intermediate values of *k* (3 ≥ *k* ≥ 5). On the other hand, when *k* is large enough, adding shape information greatly increases the dimensionality of the feature space, and the gain from adding shape information does not offset the cost of the curse of dimensionality. Thus, in this situation, the di-mismatch+shape kernel only leads to marginal improvement or in some cases even decreases the prediction performance.

We also looked at the *R*^2^ improvements for different TF families between the di-mismatch model and the di-mismatch+shape model (Figure 5, Supplementary Figure 4). For instance, in Figure 5 where *k* = 4 and *m* = 2, we observed that similar to Figure 3, adding shape features led to substantial improvements in *R*^2^ values for various zinc fingers, bHLH and HTH TFs. In addition, we found that combining shape features into the di-mismatch kernel contributed to the prediction improvements for homeodomain TFs. This observation is consistent with previous reports that specific homeodomain residues play key roles in recognizing DNA binding sites through shape readout (Dror *et al.*, 2014). For T-box TFs, since T-box proteins can bind to the DNA not only in a monomeric manner but also in dimeric combinations with various spacing and orientation patterns (Jolma *et al.*, 2013), our results suggest that the di-mismatch+shape model might help in recognizing the flexibility in the event of combinational TF bindings. Generally, our results seem to indicate that the di-mismatch kernel better describes binding sites with spacers, for instance in the center of dimeric binding targets.

**Fig. 5.**
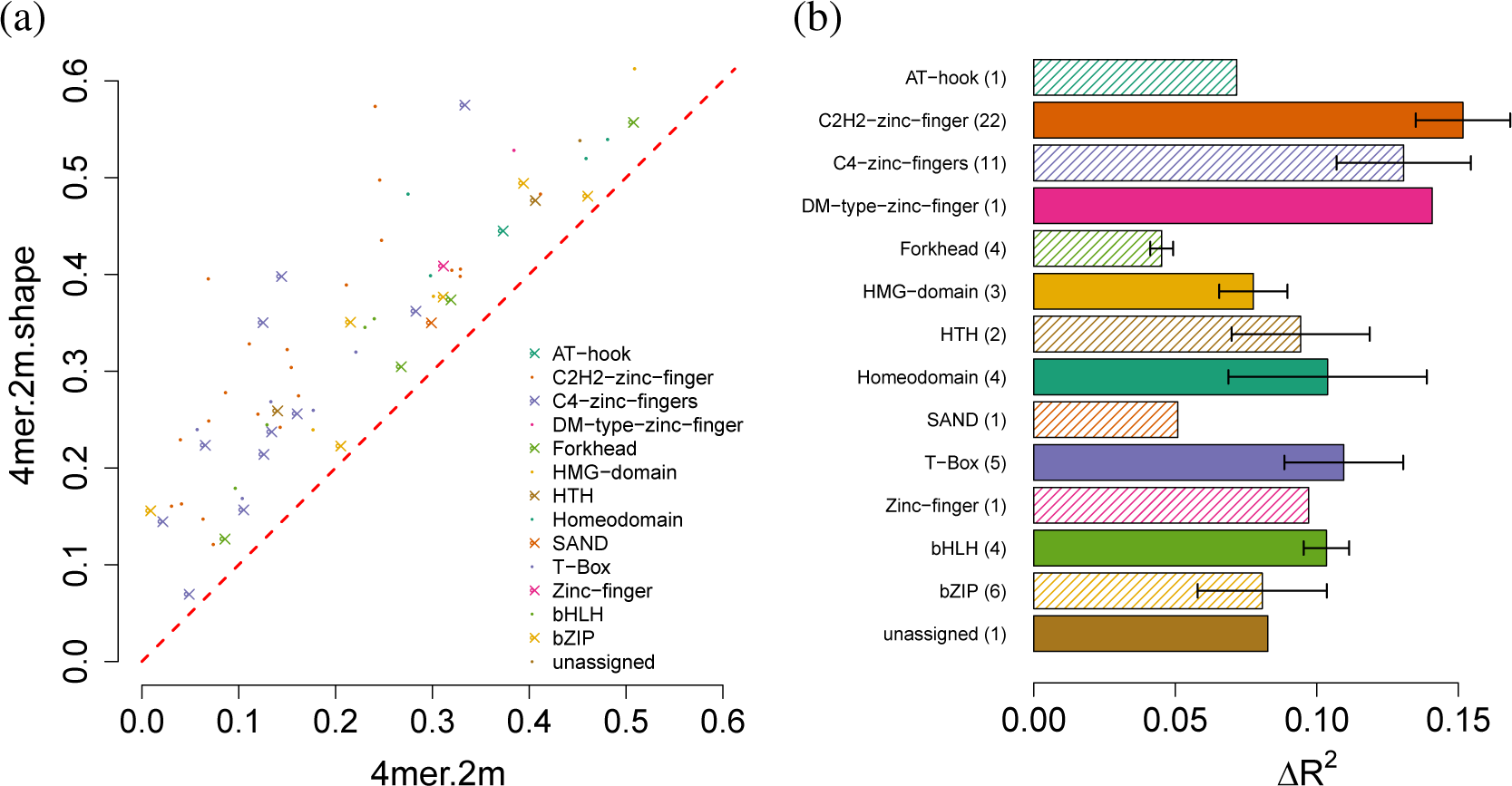
*R*^2^ performance for di-mismatch verse di-mismatch+shape model on uPBM dataset, *k* = 4 and *m* = 2.**(a)** Scatter plot of the *R*^2^ performance values between the two models. Each dot represents one TF, colored corresponding to its protein family. **(b)** Bar plot of *R*^2^ improvements for various protein families. Numbers in the parentheses are the number of DREAM5 TFs in each TF family. The *x*-axis shows the differences of *R*^2^ values between the two models. The lengths of the bars represent the mean of *R*^2^ differences and the error bars indicate the standard error of the mean.

### 4.3 The di-mismatch+shape model can accurately predict TF bindings in various experimental platforms

To investigate the extent to which our conclusions generalize beyond uPBM data, we also examined the performance of our shape-augmented models on a collection of gcPBM data for three human bHLH TFs (Myc, Mad, Max). In agreement with our observations in the mouse uPBM DREAM5 dataset, the *k*-spectrum+shape model outperformed the *k*-spectrum model for *k* < 5 for all three gcPBM datasets (Figure 6). For larger values of *k*, although the performance of the *k*-spectrum+shape model begins to drop, its *R*^2^ values are still very close to the ones for the *k*-spectrum model, for two out of three TFs. Except for the Max dataset, the best *R*^2^ performance for each the other two TF gcPBM datasets was achieved by the di-mismatch+shape model.

**Fig. 6.**
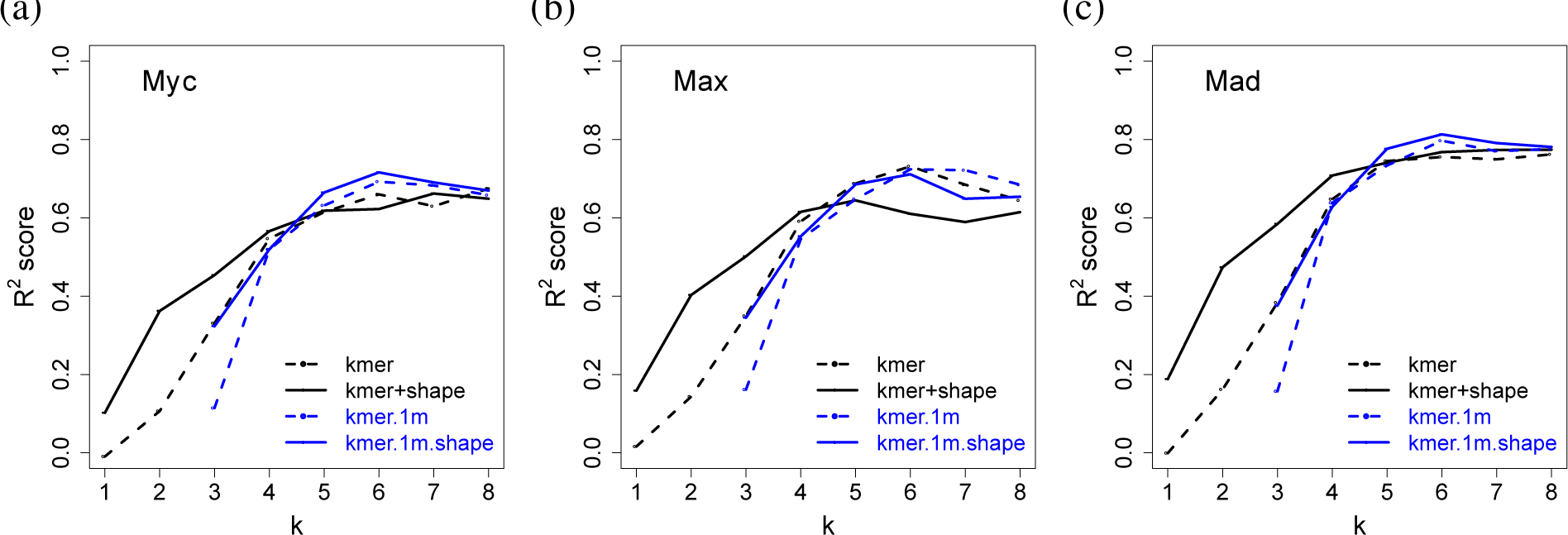
*R* 2 performance on bHLH gcPBM data. **(a)** Myc; **(b)** Max; **(c)** Mad. In each plot, dashed black line represents the performance of the *k*-spectrum model. Solid black line represents the performance of the *k*-spectrum+shape model. Dashed blue line represents the performance of the di-mismatch model. Solid blue line represents the performance of the di-mismatch+shape model. *k* = 1, …, 8.

Similarly, we observed that the di-mismatch+shape model outperformed the di-mismatch model for almost all *k* values. The benefit of adding shape information is substantial for smaller *k* values but tends to be marginal for large *k* values (*k* > 5). This might be due to the definition of the shape parameters, which require at least pentamers for the calculation of MGW.

The gcPBM dataset is of higher quality than the uPBM data, because the gcPBM data contains less positional bias and provides information on the genomic flanking regions. Therefore, we observed much higher *R*^2^ values for all the models in the human gcPBM dataset as compared to the ones in mouse DREAM5 uPBM dataset. The highest *R*^2^ value is greater than 0.8 for the Mad data. Furthermore, it has previously been shown that the flanking sequences of the 6-bp E-box core motif contribute to the binding of bHLH TFs (Gordân *et al.*, 2013). Consistent with this observation, we found that longer *k*-mers (*k* ≥ 6) in both *k*-spectrum+shape and di-mismatch+shape models continue to yield high *R*^2^ prediction accuracies for all three bHLH TFs.

### 4.4 Using DNA shape information improves the ability to distinguish between Scr and Antp binding sites

In addition to testing our shape-augmented models in a regression setting on PBM datasets, we also investigated the performance of our kernels in a classification setting to distinguish motif binding sites between two homologous Hox proteins in presence of the shared cofactor Exd. This is considered a challenging task, because the two Hox proteins, Scr and Antp, are known to bind to a similar consensus motif with subtle differences in the binding sites. Abe *et al.* (2015) previously reported that Scr and Antp recognize distinct DNA shape readout. Therefore, effectively decoding DNA shape differences is crucial to the success of distinguishing the differential binding events between Scr-Exd and Antp-Exd heterodimers.

As seen in Table 1, the *k*-spectrum+shape models consistently generated higher AUROC scores for all *k* values. In addition, the di-mismatch+shape models benefited from the inclusion of shape information and performed better than the di-mismatch models in most of the experiments when 3 ≥ *k* ≥ 7. The highest prediction AUROC score of 0.9885 was achieved by the di-mismatch+shape model with parameters *k* = 6 and *m* = 3. Therefore, our results demonstrate that with the assistance of DNA shape information we can more accurately distinguish between the binding sites of Exd-Scr and Exd-Antp heterodimers.

## 5 Discussion

Recent studies on DNA shape readout suggest that the double-helix DNA shape features play an important role in DNA binding site recognition (Rohs *et al.*, 2009). Several computational models have been developed to incorporate DNA shape information into sequence motif models and to use shape to improve the prediction accuracy of TF-DNA binding models (Zhou *et al.*, 2015; Yang *et al.*, 2014; Dror *et al.*, 2014; Mathelier *et al.*, 2016).

In this study, we present two shape-augmented models. The first one, the *k*-spectrum+shape model, is built on the classic *k*-spectrum model. The second is the di-mismatch+shape model which extends the recently developed di-mismatch model. Unlike existing sequence+shape models (Zhou *et al.*, 2015; Mathelier *et al.*, 2016), our new shape-augmented models are compositional, that is, they do not require the alignment of sequences at motif binding sites. The compositional model is better than a positional model because a compositional approach allows us to perform alignment-free modeling on all available sequences. For some TFs, we might not have a pre-defined motif model to use in creating an alignment. Furthermore, even with a well defined TF motif, there might be some sites that are transiently bound without an obvious sequence motif. Such DNA sequence might still have shape similarities that are transiently recognized (Dror *et al.*, 2016) and therefore could be recognized by our models.

Previous methods treat shape features and sequence features independently, by defining the feature vector as the concatenation of sequence features and shape features (Zhou *et al.*, 2015). Since shape features are derived from sequence information, simply adding sequence and shape information introduces redundancies in the feature space and may not be desirable. Our di-mismatch+shape shape kernel defines similarity between shape features conditioning on sequence similarity, thereby explicitly representing dependences between sequence and shape features.

Adding shape features inevitably increases the dimensionality of the feature space. To combat the curse of dimensionality, we employed a straightforward feature selection procedure. In addition, our SVM/SVR parameter *C* implicitly controls the kernel space dimension. We expect that more sophisticated feature selection approaches, such as incremental selection or regularizers like LASSO (Tibshirani, 1996) or elastic net (Zou and Hastie, 2005) could further improve our models in high-dimensional situations.

All the kernels discussed in this study encode sequence (and shape) information into vectors of features and then use linear kernels (scalar product) as the similarity score. Another possibility is to use a Gaussian (RBF) kernel. The RBF kernel embeds the data into (a finite subspace of) an infinite dimensional feature space, thus allowing efficient mapping to a high-dimensional, implicit feature space. Hence the RBF kernel might provide an alternate solution for the high-dimensionality issues in our shape-augmented models.

## Acknowledgments

This work has been supported by National Institutes of Health award R01 GM106056 (to R.R. and W.S.N.). R.R. is an Alfred P. Sloan Research Fellow.

